# Standardized droplet preamplification method for downstream circulating cell-free DNA analysis

**DOI:** 10.1101/2025.02.09.637351

**Authors:** Colin Skeen, Erica D. Pratt

## Abstract

Circulating cell-free DNA (ccfDNA) can be found in blood and other biofluids and is a minimally invasive biomarker for several pathological processes. As tumors become more invasive, an increasing amount of circulating tumor DNA (ctDNA) is also shed into the peripheral circulation. Combined analysis of ccfDNA and ctDNA has demonstrated prognostic and predictive value in metastatic disease. However, localized tumors shed significantly less ccfDNA/ctDNA and accurate detection remains a technical challenge. To overcome this barrier, droplet preamplification has been used to perform robust multiplexed analysis of low input samples. To reduce false positives, it is essential to use a high-fidelity polymerase with 3’-5’ exonuclease activity. However, attempts to combine high-fidelity polymerases with commercial droplet digital chemistries have had limited success. There is also no standardized method for efficient amplicon recovery from droplets. In this work, we present a method to reliably stabilize emulsions and recover preamplified template. We systematically compared our protocol with different destabilization methods and found an average 41% improvement in recovery efficiency. We anticipate this standardized method will increase the consistency and reproducibility of ccfDNA/ctDNA analyses. This technique could be readily translated to other low-input or low-biomass samples, such as urine, saliva, or archived biopsy specimens.

## Introduction

Circulating cell-free DNA (ccfDNA) is heavily fragmented double-stranded DNA present in low concentrations in the blood, urine, and other body fluids. In the bloodstream, ccfDNA is primarily released by hematopoietic cells undergoing apoptosis or necrosis. Since the half-life of ccfDNA is under three hours, persistence at elevated levels can be an indicator of ongoing cell death. Total ccfDNA concentration increases in several pathological processes, such as surgical trauma[1], infection[2], myocardial infarction[3], and cancer[4–9]. In cancer patients, a fraction of ccfDNA is tumor-derived and generally increases with disease stage[10]. This circulating tumor DNA (ctDNA) reflects the genetic heterogeneity of the primary tumor and any metastatic lesions. A growing body of literature has demonstrated the prognostic and predictive value of ctDNA in many high-incidence and high-mortality metastatic cancers, such as breast, colorectal, gastroesophageal, and pancreatic cancer. ccfDNA and ctDNA can serve as minimally invasive ‘liquid’ biopsies that provide continuous information at every stage of patient treatment.

However, amplification and detection of circulating DNA in early-stage disease remains challenging. Compared to metastatic disease, both overall ccfDNA concentration and ctDNA copy number are significantly reduced[11,12]. Additionally, certain high-mortality cancers (e.g. glioma, pancreatic) shed much less ctDNA into the circulation than others[13]. Multiple studies have confirmed that next-generation sequencing (NGS), digital droplet PCR (ddPCR), and other PCR-reliant platforms have reduced sensitivity and increased intra-assay variability under low input conditions[14– 17]. Stochastic amplification at low concentrations can skew the distribution of wild-type and tumor-derived sequences[18,19]. Sequence-dependent polymerase bias and errors can also prevent detection of rare mutations. The combination of amplification bias and PCR-induced errors can obscure rare, but clinically meaningful, mutations.

One example is the CXGG ‘poison’ motif present in *KRAS* (codon 12) and *TP53* (codon 248 and 282) mutations. These motifs are hypothesized to reduce polymerase processivity and impede amplification of mutant sequences[20,21]. Improving low-input assay performance could enable earlier and more effective interventions.

Several pre-PCR enrichment strategies have been developed to improve ctDNA detection from low input samples. Qualitative methods selectively amplify mutant fragments[22,23], block wild-type ccfDNA amplification [24,25], or remove wild-type ccfDNA entirely using nucleases[26,27], CRISPR/Cas9[28,29], or other site-selective cleavage proteins[30,31]. However, these approaches cannot capture quantitative changes in ccfDNA that have been associated with progression and response to therapy. Targeted preamplification of specific loci increases total DNA input while preserving both wild-type and tumor-derived sequences[32–34]. We and others have developed specialized protocols to perform targeted enrichment in bulk PCR reactions[33] and in droplets[35–39]. Multiple studies have reported that using a high-fidelity polymerase with 3’-5’ exonuclease activity reduces misincorporation errors and improves assay performance[33,36]. Okada et al. recently demonstrated that droplet preamplification significantly improved ctDNA detection and more accurately predicted the presence of *KRAS* mutant tumors (AUC 0.855 – 0.903) compared to standard ddPCR (AUC 0.430 – 0.528) [40]. Droplet preamplification has been used for ultrasensitive profiling of early-stage cancer and precancerous lesions using plasma, fine needle aspirates, and formalin-fixed paraffin-embedded tissues[35–38,41]. However, there is no standardized protocol for ensuring specific and robust pre-PCR enrichment, or efficient amplicon recovery from droplets. The lack of standardization makes it difficult to interpret results from different studies and laboratories.

An ideal pre-PCR enrichment protocol would enable both 1) unbiased amplification of multiple sequences while minimizing PCR-induced errors and 2) efficient template recovery for downstream analysis using NGS or ddPCR. Based on these criteria, the primary objectives of this study were to identify a method to stably emulsify high-fidelity PCR master mixes using Bio-Rad™ droplet digital chemistry and to reliably extract preamplified template for downstream analysis. Four different droplet destabilization strategies were evaluated based on DNA yield, DNA integrity, and protocol repeatability. Protocol performance was evaluated using contrived samples and quantified using ddPCR.

## Materials & methods

### Reagents and equipment

The dMIQE checklist and experimental details can be found in Supplementary Materials. Primers and probes were used as previously described[36].

### Reference materials

125 bp double-stranded gBlock gene fragments were used to simulate fragmented ccfDNA [42]. The positive control gBlock contained a *KRAS* G12D mutation (GAT) while the negative control contained the wild-type sequence (GGT). All gBlocks were synthesized by IDT (IA, USA). *KRAS* wild-type and *KRAS* G12D stock solutions were mixed at a 50/50 ratio to create a duplex reference standard (KRAS^WT/G12D^). Preparation details are available in Supplementary Materials.

### Droplet preamplification

Duplex preamplification reactions were prepared using the KRAS^WT/G12D^. Droplets were stabilized by adding 1 – 7 µL of 2x ddPCR Buffer Control for Probes (Bio-Rad, CA, USA) to a total reaction volume of 22 µL. Droplets were generated directly into thin-walled PCR tube strips using the Bio-Rad AutoDG Droplet Digital PCR System.

Dropletized reactions were thermal cycled with the following PCR conditions: 3 min at 98°C, 9 cycles at 98°C for 20s, 66.5°C for 20s and 72°C for 3 min, followed by 72°C for 3 min. All steps were performed using a Bio-Rad T100 thermal cycler. Master mix details are available in Table S1.3.

### Droplet destabilization

Digital preamplification reactions were set up as described above, but the polymerase was replaced with an equivalent volume of water. To accurately calculate the initial number of copies in each sample, an equivalent volume of KRAS^WT/G12D^ standard was analyzed directly using ddPCR (n = 3). Thermal cycled droplets underwent three different methods of droplet disruption (chemical, physical, and thermal). For chemical and thermal protocols, the aqueous phase containing amplified template was processed using a MinElute PCR Purification kit (Qiagen, Hilden, Germany) following a modified version of the vendor’s instructions. pH indicator was not added to Buffer PB and an additional 5-minute incubation at 35°C was added prior to elution. The final elution volume was 10 µL. The kwpower function in R package MultNonParam was used to calculate sample size for each protocol (n = 24). Based on our analysis, we had 80% power to detect differences in destabilization efficiency across groups.

#### Chemical

The chloroform extraction protocol outlined in the Bio-Rad ddPCR application guide was used as a positive control [43]. Briefly, excess droplet generation oil was removed, and samples were diluted using 20 µL of TE Buffer. Next, 70 µL chloroform was added to all samples. After vortexing at max speed for 1 minute, samples were centrifuged at 15,500 x g for 10 min. The aqueous phase containing template was collected for purification.

#### Physical

Physical droplet disruption was performed using the GeneJet Purification kit (Fisher Scientific, MA, USA) as previously described[39]. 40 µL Binding Buffer and 40 µL isopropanol were added to each sample. Samples were vortexed at top speed for 1 minute, transferred to silica columns, and centrifuged for 1 minute at 16,000 x g. 700 µL Wash Buffer was added, the sample was centrifuged, and the flow-through was discarded. Columns were centrifuged again for 1 minute. 20 µL Elution Buffer was added, and columns were centrifuged for 1 minute. The physical disruption protocol simultaneously purified the template and was analyzed directly.

#### Thermal (Thermal^a^)

PCR tube strips were submerged in liquid nitrogen for 1 minute, then thawed to room temperature as previously described [44]. The aqueous phase was collected, and the total volume was recorded prior to purification.

#### Modified Thermal (Thermal^b^)

Based on observed results using previously reported methods, we created a revised thermal protocol. Droplets were diluted using 40 µL of TE Buffer and vortexed at max speed for 10 min. Samples were centrifuged at 2,000 x g for 1 min, snap frozen using liquid nitrogen for 1 min, and thawed at RT. This process was repeated and then the aqueous phase containing template was collected for purification. DNA integrity following thermal disruption was analyzed using TapeStation cfDNA ScreenTape (Agilent Technologies, CA, USA).

### ddPCR workflow

All reagents were thawed to room temperature for 30 minutes before preparing ddPCR reactions (Table S1.4). Droplets were generated in semi-skirted plates and transferred to a C1000 Touch™ thermal cycler. Reactions were thermal cycled with the following PCR conditions: 10 minutes at 95°C; 40 cycles of 30s at 94°C, 60s at 61°C; 10 min. at 98°C, and a final hold at 12°C. The ramp rate was 2°C/s. Once the thermal cycler lid volume was below 50°C, plates were removed and analyzed using the QX600 Droplet Reader. Samples with more than 10,000 intact droplets were analyzed using QX Manager 2.0 (Bio-Rad). Gating thresholds were set manually based on the droplet amplitude of non-template controls (NTCs) and applied uniformly to all experimental samples.

### Performance metrics

DNA preamplification efficiency was defined as follows:

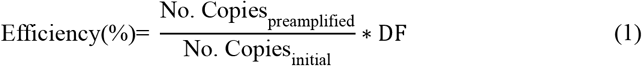

Where No. Copiesinitial was defined as the average measured concentration of the unamplified sample using ddPCR (n = 3). The dilution factor (DF) was defined as the ratio of the volume of template analyzed and total template volume.

The percentage of DNA recovered was defined as follows:

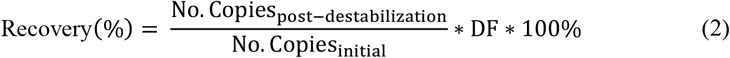

### Statistical analysis

Unless otherwise noted, data is reported as median and standard deviation (SD) using the form median (SD). To compare statistical differences between multiple samples, the non-parametric Kruskal-Wallis test was used. Results significant by Kruskal-Wallis were followed by a Dunn’s post-test with Benjamini-Hochberg correction for multiple comparisons. Comparisons of two groups were made using a non-parametric two-tailed Wilcoxon-Mann-Whitney test. Homogeneity of variances was tested using the non-parametric Fligner-Killeen test. For all hypothesis testing, the minimum level of statistical significance (p-value) was 0.05. R version 4.1.3 (The R Foundation for Statistical Computing, Vienna, Austria) was used for data analysis and figure construction.

## Results & discussion

### Stabilization of droplets containing high-fidelity polymerase reagents

Prior reports using non-proprietary PCR mixes or additives with Bio-Rad™ droplet digital chemistries have reported stability issues including polydispersity, coalescence, or failure to generate droplets entirely[38,45,46]. Adding a small volume of ddPCR Supermix was reported to stabilize LAMP reactions[46]. However, the polymerase present in ddPCR Supermix has a high false-positive rate (∼0.10%) and is unsuitable for preamplification of low input samples[38]. ddPCR Buffer Control has a viscosity similar to ddPCR Supermix but does not contain a polymerase. It is sold as a blank control and is fully compatible with the Bio-Rad Automated Droplet Generator (AutoDG). Based on these findings, we tested whether Buffer Control could be used to stabilize emulsions containing high-fidelity PCR reagents. All experiments used Q5 High-Fidelity polymerase, which has been previously shown to enable efficient and accurate preamplification of low input ccfDNA samples[33,36,37].

To evaluate the impact of Buffer Control on droplet stability, a titration series experiment was conducted (1 µL – 7 µL) and droplets were thermal cycled and evaluated by visual inspection. Without Buffer Control, droplets were successfully generated but would coalesce after thermal cycling (Fig 1A). A discrete droplet layer was observed starting at 2 µL and droplet homogeneity appeared to increase with Buffer Control added. To identify the minimum required volume of Buffer Control, we measured the preamplification efficiency across a narrower range (1.5 – 4.5 µL). Each reaction contained 2,850 copies of the KRAS^WT/G12D^ reference standard. Amplification efficiency increased with Buffer Control added until 3.5 µL, after which there was no statistically significant difference (Fig 1B). Two-dimensional cluster plots of the duplex reaction showed clear separation of the two molecular targets (Fig 1C). Non-template and wild-type only controls showed no observable increase in false positives compared to conventional ddPCR (Fig S3, Fig S4). We selected 4 µL of Buffer Control as the optimized volume to ensure small variations in sample quality would not impact assay performance. The median amplification ratio for the 4 µL group was 60.15 (14.27%) which matches our prior results[36]. The preamplification efficiency of *KRAS* WT and G12D were similar (p = 0.297; Wilcoxon-Mann-Whitney), suggesting no significant amplification bias (Fig 1D). Based on these results, we used the optimized preamplification protocol for subsequent experiments.

**Figure 1.**
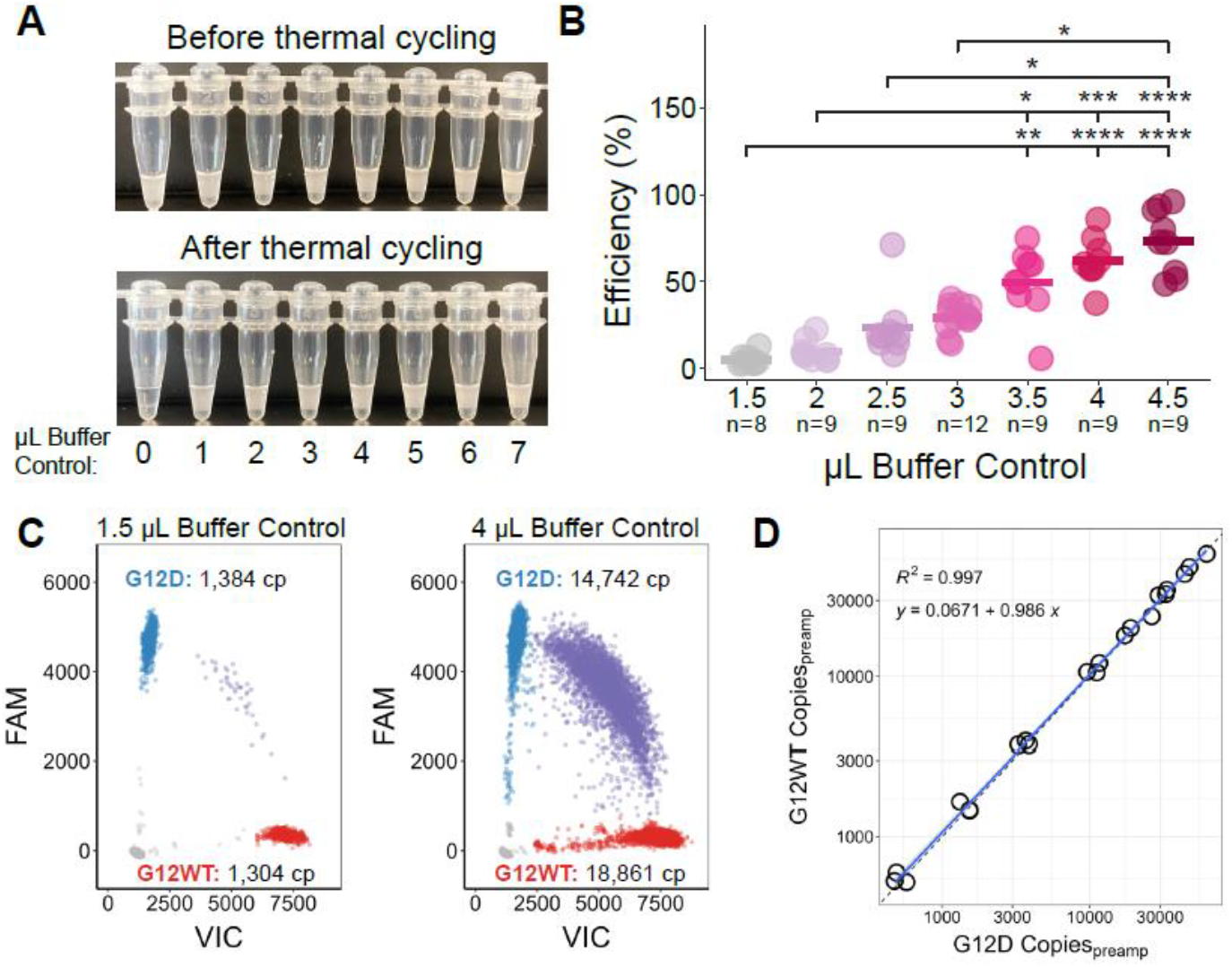
Optimization of droplet preamplification reaction. A) Representative droplets generated using Q5 high-fidelity polymerase and varying amounts of Bio-Rad Buffer Control for Probes before (top) and after (bottom) thermal cycling. B) Efficiency of PCR preamplification as a function of Buffer Control added to the reaction mix (Kruskal-Wallis). C) Representative 2D plot for pessimal (left) and optimal (right) reaction mixes. D) Comparison of total number of amplified copies for *KRAS* WT and *KRAS* G12D in a duplex reaction. ****: p<0.0001; ***: p<0.001; **: p<0.01; *: p<0.05.

### Comparison of droplet destabilization methods

For accurate detection of low abundance ctDNA in early-stage cancer, efficient recovery of amplicons from droplets is critical. Bio-Rad recommends a chloroform destabilization protocol to extract DNA from droplets [43]. More recently, physical [39] and thermal [44] destabilization protocols have reported similar DNA recoveries with fewer processing steps. Chloroform displaces surfactants from the oil-water droplet interface, inducing coalescence. However, chemical destabilizers are partially soluble in the recovered aqueous layer and can reduce the efficiency of downstream PCR reactions [47]. Thermal destabilization methods use liquid nitrogen freeze-thaws to expand the inner aqueous layer, causing the interfacing surfactant to become sparser and decreasing droplet stability. Physical destabilization requires passing the droplets through a DNA spin column with centrifugation steps to physically disrupt the droplets. Both methods reduce opportunities for cross-contamination and sample loss.

To determine which method worked best with our optimized protocol, we compared the number of copies present in a reference standard of known concentration before and after dropletization and destabilization (Fig 2A). Preamplification reactions were set up as previously described, but the polymerase was replaced with an equivalent volume of water. This yielded stabilized DNA-containing droplets that could be thermal cycled, but no amplification would take place. In our hands, the Bio-Rad recommended chemical protocol outperformed both physical and thermal protocols (Table 1). Our results are in line with the previously reported recoveries for chemical (45%) and physical (36%) protocols [44]. However, our measured recovery for thermal destabilization (Thermal^a^) was significantly lower than previously reported (27.4% versus 68%). A recent study using droplet preamplification to improve NGS-based ccfDNA analysis also reported lower recovery when using thermal destabilization[39]. Our master mix modifications to emulsify high-fidelity reagents could increase droplet stability, which might explain this difference in results.

**Table 1.** Efficiency and variability of four different droplet destabilization methods. ^1^ standard deviation ^2^ coefficient of variation

| Method | Recovery (%) | SD<sup>1</sup> (%) | CV<sup>2</sup> (%) |
| --- | --- | --- | --- |
| Chemical | 51.2 | 9.0% | 17.6 |
| Physical | 46.6 | 16.7% | 35.8 |
| Thermal<sup>a</sup> | 30.4 | 12.86% | 42.3 |
| Thermal<sup>b</sup> (our protocol) | 68.2 | 14.87% | 21.8 |

**Figure 2.**
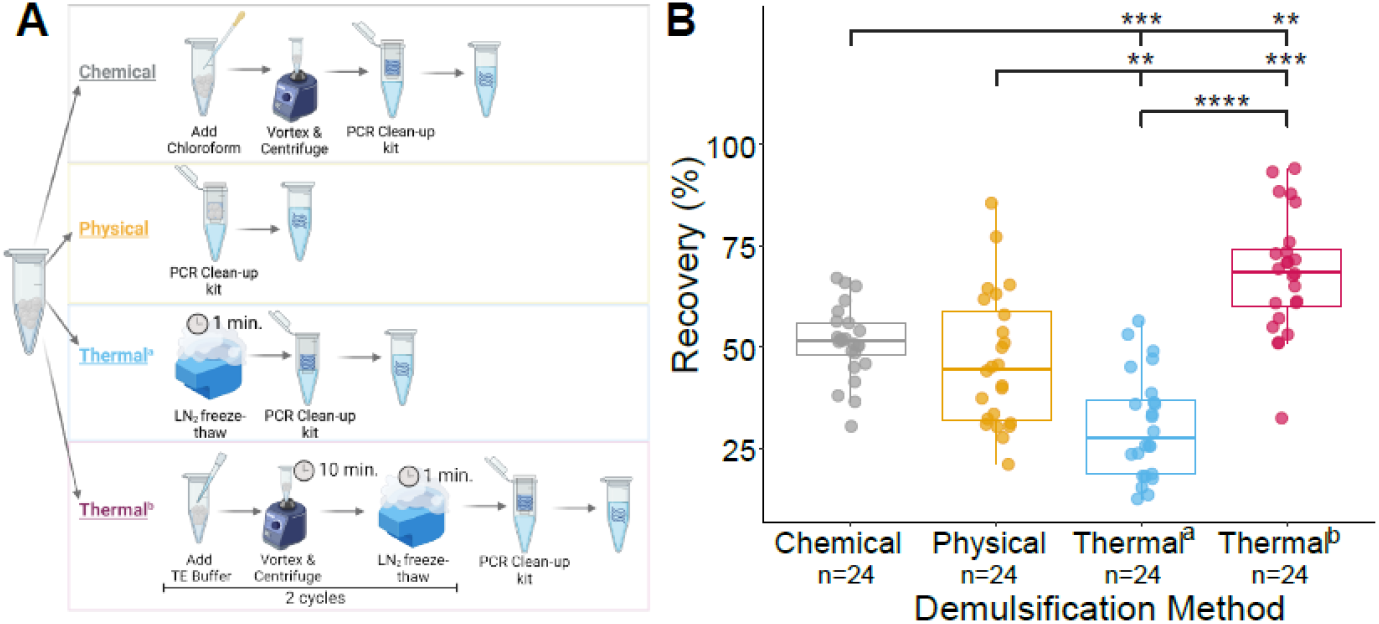
Evaluation of droplet destabilization methods. A) Experimental design for evaluating each protocol. B) The proportion of input template recovered using chemical, physical, and thermal protocols (Kruskal Wallis; Dunn’s post-hoc). ****: p<0.0001; ***: p<0.001; **: p<0.01; *: p<0.05.

We then created a hybrid protocol containing elements of the chemical and thermal protocols, but only using TE buffer as a diluent. We found that the new thermal protocol (Thermal^b^) had the highest percent recovery compared to prior methods (p <0.00001; Kruskal-Wallis) (Fig 2B). The introduction of a dilution and vortexing step did not affect assay repeatability, as variance in DNA recovery was not significantly different across the four methods (p < 0.05; Fligner-Killeen). We also tested variables such as centrifugation speed, and altered the order of the dilution, vortexing, and freeze-thaw steps. None of these modifications improved DNA recovery (Fig S5).

### Evaluation of DNA integrity

A concern with thermal methods is that ccfDNA can be damaged by multiple freeze-thaw cycles [48,49]. To evaluate whether the two liquid nitrogen steps in the Thermal^b^ protocol impacted DNA quality, we analyzed gBlock samples before and after destabilization using capillary electrophoresis. The original 125 bp fragment and 95 bp amplicon were clearly distinguishable with no evidence of additional fragmentation or degradation (Fig 3, Fig S6).

**Figure 3:**
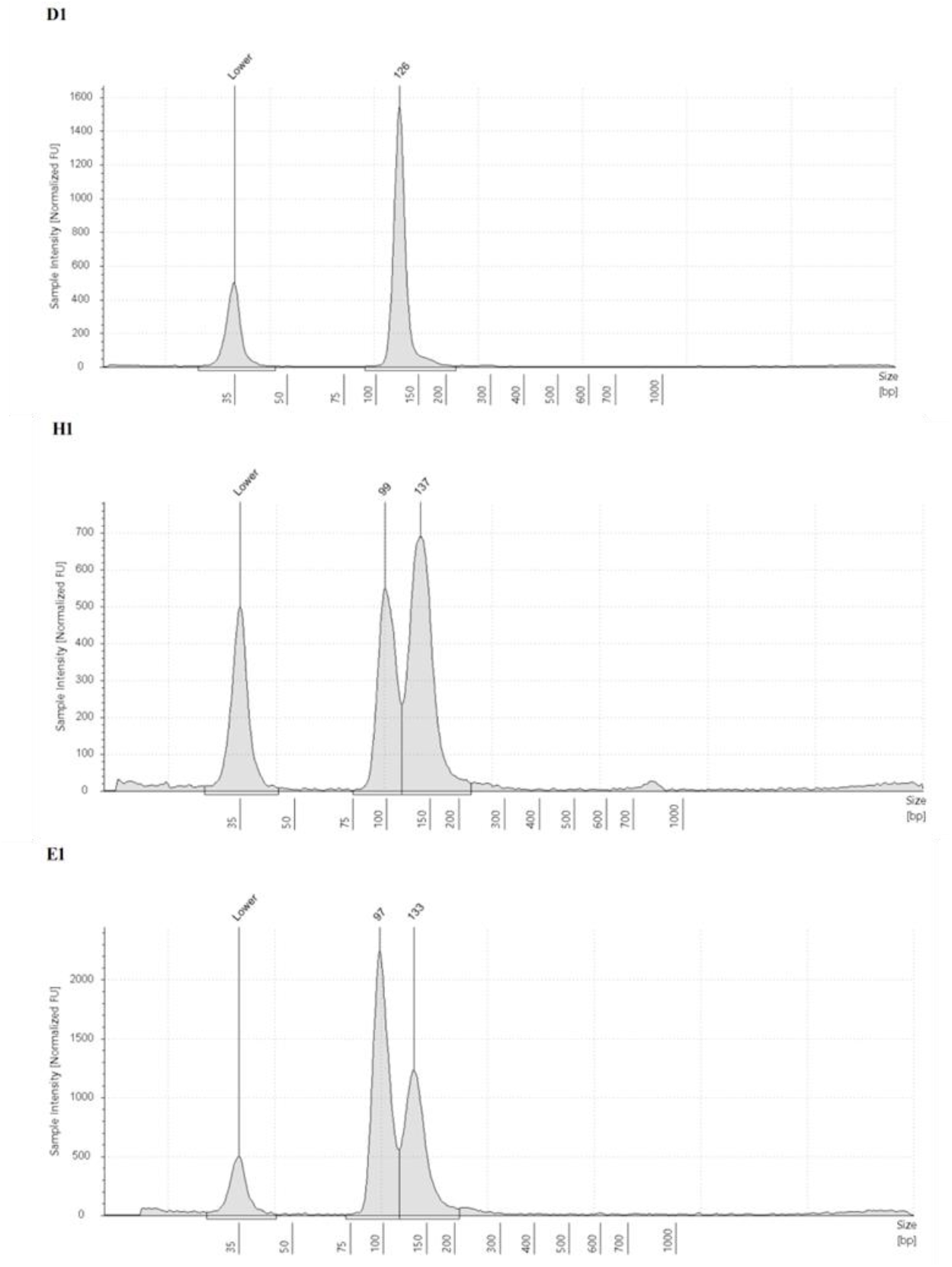
Representative Tapestation electropherograms of gBlocks pre- and post-droplet preamplification. D1: 125 bp KRAS gBlock before preamplification. H1 & E1: Samples taken after thermal destabilization (protocol Thermal^b^). Only two peaks are visible, the original unamplified gBlock and the expected 95 bp amplicon. *Lower* is the lower marker used to align ladder data with samples of unknown size.

## Conclusion

The goal of this study was to develop a unified protocol for ccfDNA preamplification using Bio-Rad droplet digital chemistry. To stabilize high-fidelity polymerase reagents, we leveraged existing Bio-Rad reagents. This approach is fully compatible with automated droplet-generator systems and minimizes any chance of contamination or line fouling. To recover preamplified template, we developed a destabilization protocol with an efficiency of 68.6%. Using gBlocks as contrived ccfDNA samples allowed for standardized evaluation of our method and previously published techniques. Our approach outperformed chemical, physical, and thermal methods by 33%, 53%, and 150%, respectively.

## Acknowledgements

We thank Dr. Susan Tsai, Dr. Poojitha Sitaram, and Julia Sexton for helpful discussion, suggestions, and early technical assistance.

## Future Perspective

Droplet preamplification can resolve subsampling effects that are a key barrier preventing accurate quantification and analysis of low input plasma samples. This allows for repeated testing of each plasma sample without exhausting the original specimen, improving assay robustness and enabling analyses using orthogonal methods. Future studies could implement synthetic spike-in controls[49] to quantify amplification efficiency and DNA recovery from droplets with even higher accuracy. While this study leverages existing commercial ddPCR reagents, this approach could be readily adapted to other droplet microfluidics platforms. This pre-PCR enrichment method maximizes the amount of information that can be obtained in a single blood draw.

