## Supplementary Information for "Standardized droplet preamplification method for downstream circulating cell-free DNA analysis"

*Table of Contents*


| | |
|---|---|
| *dMIQE information* | 2 – 7 |
| *Supplementary Methods* | 8 |
| *Supplementary Figures* | 9 – 12 |
| *References* | 13 |

### S1. Minimum Information for Publication of Digital PCR Experiments (dMIQE)

#### Table S1.1: dMIQE checklist for authors, reviewers, and editors

| ITEM TO CHECK | PROVIDED | COMMENT |
|---|---|---|
| **1. SPECIMEN** | | |
| Detailed description of specimen type and numbers | Y | Materials & methods, S2 |
| Sampling procedure (including time to storage) | N | NA |
| Sample aliquoting, storage conditions and duration | Y | Materials & methods, S2 |
| **2. NUCLEIC ACID EXTRACTION** | **NA** | |
| Description of extraction method including amount of sample processed | N | |
| Volume of solvent used to elute/resuspend extract | N | |
| Number of extraction replicates | N | |
| Extraction blanks included? | N | |
| **3. NUCLEIC ACID ASSESSMENT AND STORAGE** | | |
| Method to evaluate quality of nucleic acids | Y | Materials & methods, S2.2 |
| Method to evaluate quantity of nucleic acids (including molecular weight and calculations when using mass) | Y | S2.2 |
| Storage conditions: temperature, concentration, duration, buffer, aliquots | Y | S2.2 |
| Clear description of dilution steps used to prepare working DNA solution | Y | S2.2 |
| **4. NUCLEIC ACID MODIFICATION** | | |
| Template modification (digestion, sonication, pre-amplification, bisulphite, etc.) | Y | Materials & methods, Table 1.3 |
| Details of repurification following modification, if performed | Y | Materials & methods |
| **5. REVERSE TRANSCRIPTION** | **NA** | |
| cDNA priming method and concentration | | |
| One or two step protocol (include reaction details for two step) | | |
| Amount of RNA added per reaction | | |
| Detailed reaction components and conditions | | |
| Estimated copies measured with and without addition of reverse transcriptase | | |
| Manufacturer of reagents used, with catalogue and lot numbers | | |
| Storage of cDNA: temperature, concentration, duration, buffer and aliquots | | |
| **6. dPCR OLIGONUCLEOTIDES DESIGN AND TARGET INFORMATION** | | |
| Sequence accession number or official gene symbol | Y | KRAS |
| Method (software) used for design and *in-silico* verification | Y | Previously published[1] |
| Location of amplicon | Y | S2.1 |
| Amplicon length | Y | Table S1.2 |

| | | |
|---|---|---|
| Primer and probe sequences (or amplicon context sequence) | Y | Table S1.2 |
| Location and identity of any modifications | N | NA |
| Manufacturer of oligonucleotides | Y | Table S1.2, S2.1 |
| **7. dPCR PROTOCOL** | | |
| Manufacturer of dPCR instrument and instrument model | Y | Materials & methods |
| Buffer/kit manufacturer with catalogue and lot number | Y | Table S1.5 |
| Primer and probe concentrations | Y | Table S1.3, Table S1.4 |
| Pre-reaction volume and composition (including amount of template and if restriction enzyme added) | N | NA |
| Template treatment (initial heating or chemical denaturation) | N | NA |
| Polymerase identity and concentration, $Mg^{++}$ and dNTP concentrations | Y | Table S1.3, Table S1.4 |
| Complete thermocycling parameters | Y | Materials & methods |
| **8. ASSAY VALIDATION** | | |
| Details of optimisation performed | Y | Results & discussion, Fig S3-S5 |
| Analytical specificity (vs. related sequences) and limit of blank | N | NA |
| Analytical sensitivity/limit of detection, and how this was evaluated | N | NA |
| Testing for inhibitors (from biological matrix/extraction) | N | NA |
| **9. DATA ANALYSIS** | | |
| Description of dPCR experimental design | Y | Materials & methods |
| Comprehensive details of negative and positive controls (whether applied for quality control or for estimation of error) | Y | Materials & methods |
| Partition classification method (thresholding) | Y | Materials & methods |
| Examples of positive and negative experimental results (including fluorescence plots in Supplemental Materials) | Y | Fig 1, Fig S6 |
| Description of technical replication | Y | Materials & methods, Results & discussion |
| Repeatability (intra-experiment variation) | Y | Results & discussion |
| Reproducibility (inter-experiment/user/lab, etc. variation) | Y | Results & discussion |
| Number of partitions measured (average and standard deviation) | N | Reactions with <10.000 partitions were not analyzed |
| Partition volume | Y | 0.795 nL (vendor reported) |
| Copies per partition (λ or equivalent) (average and standard deviation) | N | variable |
| dPCR analysis program (source, version) | Y | Materials & methods |
| Description of normalisation method | N | NA |
| Statistical methods used for analysis | Y | Materials & methods |
| Data transparency | Y | Data available upon request |

**Table S1.2: List of designed *KRAS* assays**

| Amplicon Length | Type | $T_m$(°C) | Oligonucleotide sequence | Vendor |
|---|---|---|---|---|
| 95 | F Primer | 58 | 5'-ATTATAAGGCCTGCTGAAAATGACT-3' | IDT |
| | R Primer | 58.5 | 5'-TCTGAATTAGCTGTATCGTCAAGG-3' | IDT |
| | $Probe_{WT}$ | 60.5 | 5'-**VIC**-TTGGAGCTGGTGGCGT- **MGBNFQ**-3' | Life Technologies |
| | $Probe_{G12D}$ | 56.4 | 5'-**FAM**-TGGAGCTGATGGCGT-**MGBNFQ**-3' | Life Technologies |

**Table S1.3: Mastermix for droplet preamplification method**

| Component[1] | Concentration in PCR mastermix |
|---|---|
| Buffer Control for Probes | 18% |
| Fwd Primer | 500 nM |
| Rev Primer | 500 nM |
| dNTPs | 200 µM |
| Q5 Reaction Buffer | 1x |
| Q5 High-Fidelity DNA Polymerase | 0.02 U/µl |
| Sample | 0 – 9.76 µL |

[1] Calculations for final PCR volume of 20 µL

**Table S1.4: Duplex *KRAS* ddPCR assay**

| Component[1] | Concentration in PCR mastermix |
|---|---|
| Supermix for Probes (no dUTP) | 1x |
| Fwd Primer | 900 nM |
| Rev Primer | 900 nM |
| *KRAS* WT Probe | 250 nM |
| *KRAS* G12D Probe | 250 nM |
| Sample | 0 – 7.64 µL |

[1] Calculations for final PCR volume of 20 µL

**Table S1.5: Additional reagent information**

| Component | Company | Part No. |
|---|---|---|
| Cell-free DNA ScreenTape | Agilent | 5067-5630 |
| Cell-free DNA ScreenTape Reagents | Agilent | 5067-5631 |
| ddPCR Supermix for Probes (no dUTP) | Bio-Rad | 1863025 |
| ddPCR Buffer Control for Probes | Bio-Rad | 1863052 |
| ddPCR 96-well plates | Bio-Rad | 12001925 |
| QX600TM Droplet Reader | Bio-Rad | 12013328 |
| Automated Droplet Generator | Bio-Rad | 1864101 |
| PX1 PCR Plate Sealer | Bio-Rad | 1814000 |
| C1000 Touch Thermal Cycler | Bio-Rad | 1851197 |
| T100 Thermal Cycler | Bio-Rad | 1861096 |
| Eppendorf DNA LoBind Microcentrifuge Tubes | Eppendorf | 13-698-791 |
| TE Buffer, pH 8.0, Low EDTA | Fisher Scientific | AAJ75793AP |
| Invitrogen UltraPure DNase/RNase-Free Distilled Water | Fisher Scientific | 10-977-023 |
| Benchtop Liquid Nitrogen Container | Fisher Scientific | 11-670-4B |
| GeneJET PCR Purification Kit | Fisher Scientific | FERK0701 |
| Q5® High-Fidelity DNA Polymerase | NEB | M0491 |
| Q5® Reaction Buffer Pack | NEB | B9027 |
| Deoxynucleotide (dNTP) Solution Mix | NEB | N0447 |
| 0.2 mL 12-Tube PCR Strips | USA Scientific | 1402-2400 |
| MinElute PCR Purification Kit | Qiagen | 28006 |

### S2. gBlocks

#### *S2.1 gBlock synthetic dsDNA sequences*

Flank sequence of *KRAS* obtained from dbSNP using id rs121913529. Position of codon 12 is highlighted in grey. All gBlocks were ordered from IDT.

*KRAS Wild-type*

ATTATTTTATTATAAGGCCTGCTGAAAATGACTGAATATAAACTTGTGGTAGTTGGAGCTGGT
GGCGTAGGCAAGAGTGCCTTGACGATACAGCTAATTCAGAATCATTTTGTGGACGAATATG

*KRAS G12D*

ATTATTTTATTATAAGGCCTGCTGAAAATGACTGAATATAAACTTGTGGTAGTTGGAGCTGATG
GCGTAGGCAAGAGTGCCTTGACGATACAGCTAATTCAGAATCATTTTGTGGACGAATATG

#### *S2.2 Dilution of gBlocks*

gBlocks were reconstituted per vendor instructions at a theoretical concentration of 10 ng/µL or ~7.8e7 copies per microliter (cp/µL). This primary stock (1°) was diluted to 1 ng/µL or 7.8e6 cp/µL and the resulting secondary stock (2°) was quantified using the Qubit 4 Fluorometer (Thermo Scientific, MA, USA). To reach the final experimental concentrations, the 2° stock solution was serially diluted to 2.85e6 cp/µL (3°) and 2.85e3 cp/µL (4°). Stock solutions 1°– 3° were prepared in 5 µL aliquots and stored at -20°C until use. Working stock solution 4° was stored at 4°C and remade weekly. All solutions were made using TE buffer (10 mM Tris, 0.1 mM EDTA) and stored in 1.5-mL DNA LoBind tubes (Eppendorf, CT, USA). When working with low DNA concentration aliquots, we consistently observed that the copy number was 20% lower than expected. We attributed this to DNA binding to the storage container, which matches prior reports using commercial low-bind tubes[2]. Based on these data, a 20% correction was built into dilution calculations. Aliquots of the 4° stock were measured using the QX600 ddPCR system.

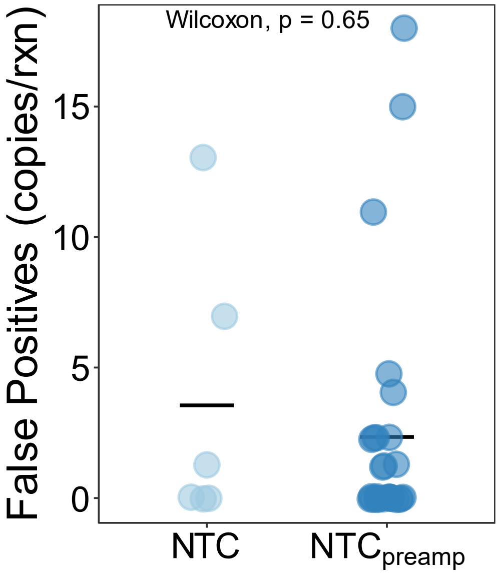


**Figure S3**: **Comparison of false positive rates between ddPCR and droplet preamplification.** The average number of false positives per reaction was 3.6 (IQR: 0-5.6) for ddPCR and 2.3 (IQR: 0-2.3) for digital preamplification (p = 0.65; Mann-Whitney-Wilcoxon). False positives were calculated for each *KRAS* target (G12D and WT). Crossbar on graph represents the mean for each group. Sample size: ddPCR (n=3); droplet preamplification (n=14).

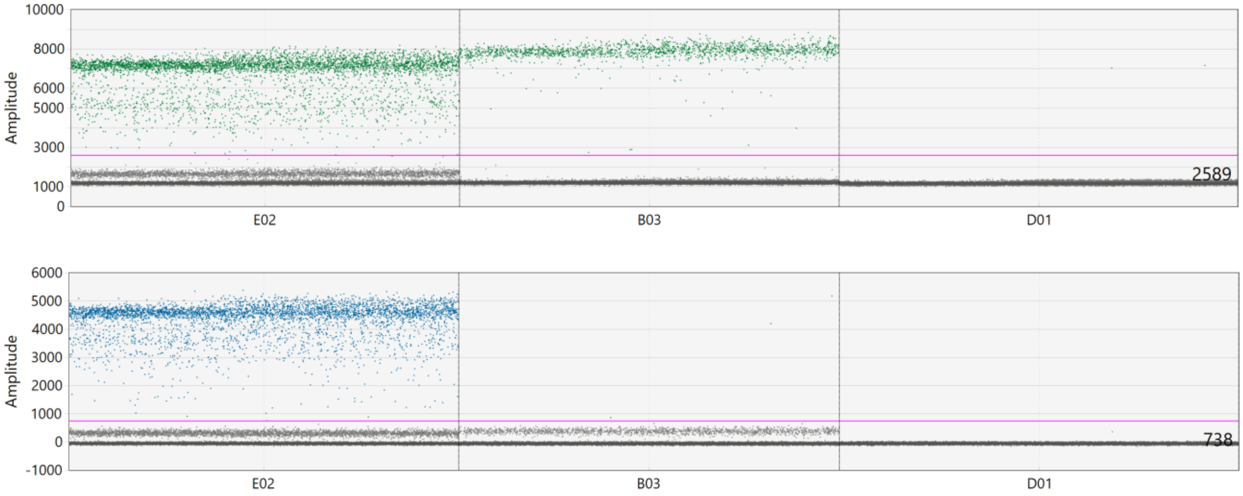


**Figure S4**: **Representative 1D amplitude view of a ddPCR run.** E02: duplex reaction for *KRAS* WT and G12D; B03: WT-only control; D01: Non-template control.

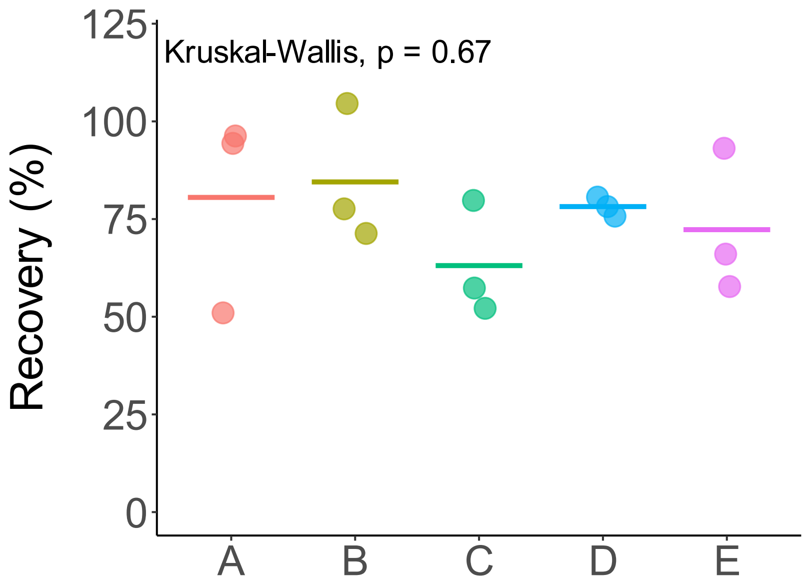


**Figure S5**: **Comparison of additional destabilization steps.** Observed recovery efficiency for five protocol changes (L-R): A) Processing both the aqueous and oil phase using a Qiagen PCR cleanup kit, B) Increasing centrifugation force from 2,000 x g to 3,217 x g, C) Removing oil phase prior to sample dilution and thermal destabilization, D) Performing the freeze-thaw step prior to sample dilution and vortexing, and E) The final Thermal$^{b}$ protocol reported in this study. Three replicates were performed for each condition.

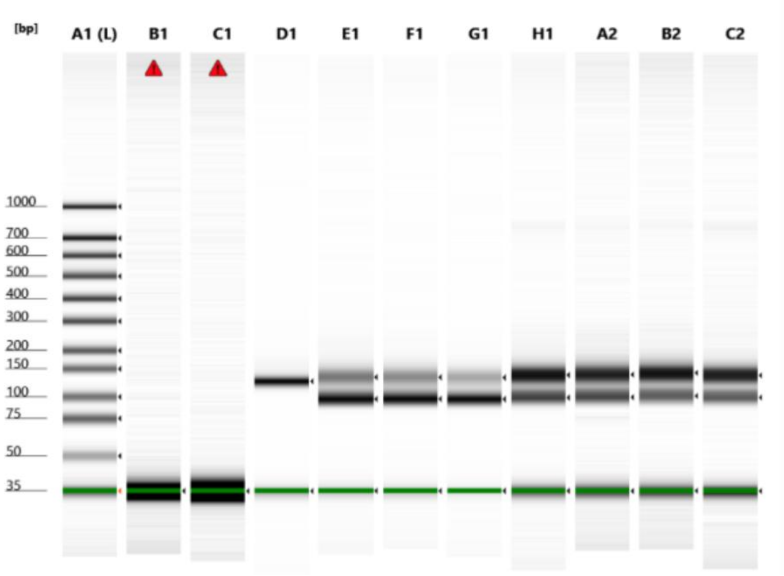


**Figure S6. Agilent TapeStation analysis of gBlock integrity before and after thermal destabilization.** A1: ladder. B1: TE buffer. C1: Droplet preamplification NTC. No signal was seen in any of the negative controls. D1: 125 bp *KRAS* gBlock. E1-C2: Seven replicates of *KRAS* gBlock after thermal recovery from droplets**.**
